# Alternative Lengthening of Telomeres is not synonymous with mutations in ATRX/DAXX

**DOI:** 10.1101/2020.05.05.076125

**Authors:** Alexandre de Nonneville, Roger R. Reddel

## Abstract

The PCAWG Consortium has recently released an unprecedented set of tumor whole genome sequence (WGS) data from 2,658 cancer patients across 38 different primary tumor sites^1^. WGS is able to document the quantity and distribution of telomeric repeats^2^. In one of the papers analyzing the PCAWG dataset, Sieverling *et al.*^3^ confirmed previous data^4^ indicating that tumors with truncating ATRX or DAXX alterations, referred to as ATRX/DAXX^trunc^, have an aberrant telomere variant repeat (TVR) distribution. By regarding these mutations, *vs.* TERT modifications (TERT^mod^; i.e. promoter mutations +/− amplifications +/− structural variations), as indicators of Alternative Lengthening of Telomeres (ALT) *vs.* telomerase, they built a random forest classifier for ALT-probability, and then associated genomic characteristics with the putative Telomere Maintenance Mechanism (TMM)^3^. However, we show here that equating ATRX/DAXX^trunc^ and TERT^mod^ with ALT and telomerase, respectively, results in TMM predictions which correlate poorly with TMM assay data. ATRX/DAXX^trunc^ mutations are heterogeneously distributed in ALT-positive (ALT+) tumors of different types, as are TERT^mod^ in telomerase-positive tumors^4^. Although these mutations are strongly associated with TMM, most tumors do not harbor them, making them an inadequate basis for building a classifier in a large-scale pan-cancer study^4–7^. Here, we provide a new analysis of the PCAWG data, based on C-circle assay (CCA)^8^ that is available for a subset of these tumors^4,9,10^. We show that the Sieverling *et al.* score overestimates the proportion of ALT associated with ATRX/DAXX^trunc^ and misclassifies ALT tumors when these mutations are absent. We also show some TVR correlate with ATRX/DAXX^trunc^ mutations, regardless of TMM. Finally, we propose a new classifier to identify ALT tumors in the PCAWG cohort.

## Poor correlation with telomere maintenance assay and previously published score

WGS data have been used as a means to assess telomere content and analyse TMM^2,5^. To our knowledge, only one previous genomic study generated an ALT-probability score based on ALT assay data. Lee et *al.*^4^ used the CCA, a reliable marker of ALT activity^8^, to determine whether the tumors were ALT+ or ALT- for 167 pancreatic neuroendocrine tumors (PaNET) and melanomas, and then applied machine learning to features including total telomeric and TVR content to develop an ALT classifier with an accuracy of 91.6%. The classifier was then applied to WGS data from 908 additional tumors, mostly from The Cancer Genome Atlas (TCGA) dataset. Of the total of 1075 tumors studied by Lee et *al.*^4^, 703 were included in the PCAWG study, and CCA data are available for 114 of these (melanoma, n=46 and PanNET, n=68^4,9,10^). A comparison of the Sieverling and Lee scores revealed a poor correlation (*r*=0.101 [95%CI 0.025-0.17], Spearman correlation; Supplementary Fig. 1). We then compared the CCA data with Sieverling’s score. The latter identified only 64.5% of CCA-positive tumors as ALT-high probability and misidentified CCA-negative ATRX/DAXX^trunc^ tumors (Figure 1). Also, the only CCA-positive TERT^mod^ tumor was identified as ALT-low probability by the Sieverling score.

**Figure 1.**
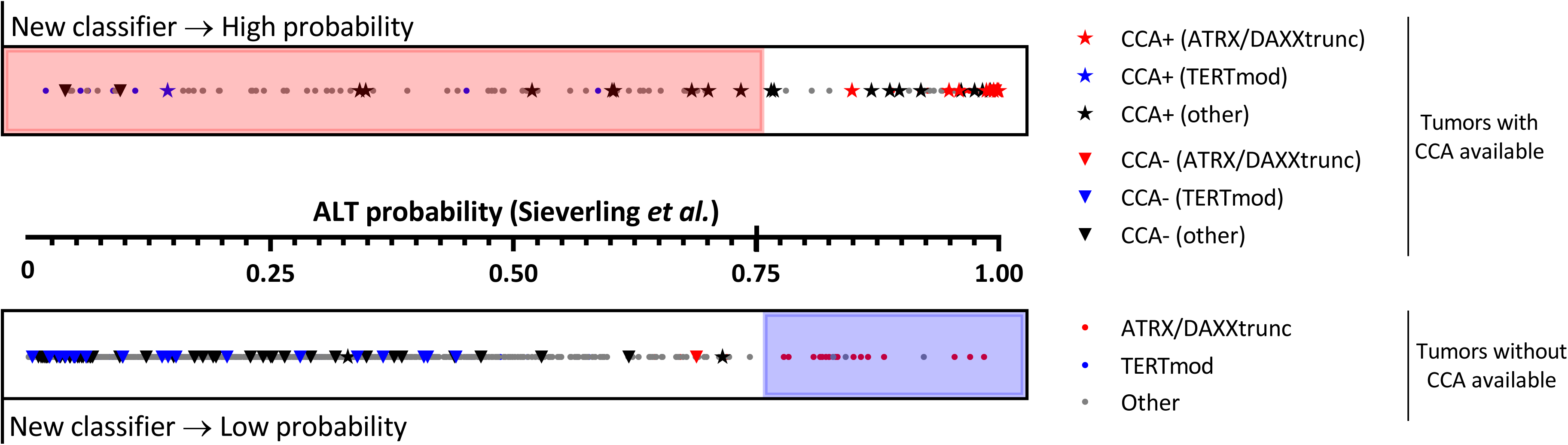
Distribution of Sieverling *et al.*^3^ ALT-probability score in 2,497 patients included in the PCAWG study, dichotomized according to our new classifier. Stars and inverted triangles represent tumor samples with CCA (C-circles assay) data available^4,9,10^. Red and blue symbols correspond to patients with ATRX/DAXX^trunc^ and TERT^mod^ alterations, respectively ^3^. Tumors misclassified by Sieverling’s score are highlighted by red (ALT-high) and blue (ALT-low) shadings.

## Specific variant distributions correlate with ATRX/DAXX mutations, regardless of the TMM

Sieverling *et al.’*s analysis of the PCAWG data claimed an association between prevalence of TTCGGG singletons and ALT, and a more pronounced enrichment of the TGAGGG TVR than previously noted. Our re-analysis showed that TTCGGG, TGAGGG, TTTGGG and TTGGGG singleton distributions are similar among ATRX/DAXX^trunc^ tumors, regardless of CCA result (Figure 2). In contrast, in CCA-positive tumors, there were statistically significant differences in TTCGGG and TGAGGG distributions between tumors with and without ATRX/DAXX^trunc^.

**Figure 2.**
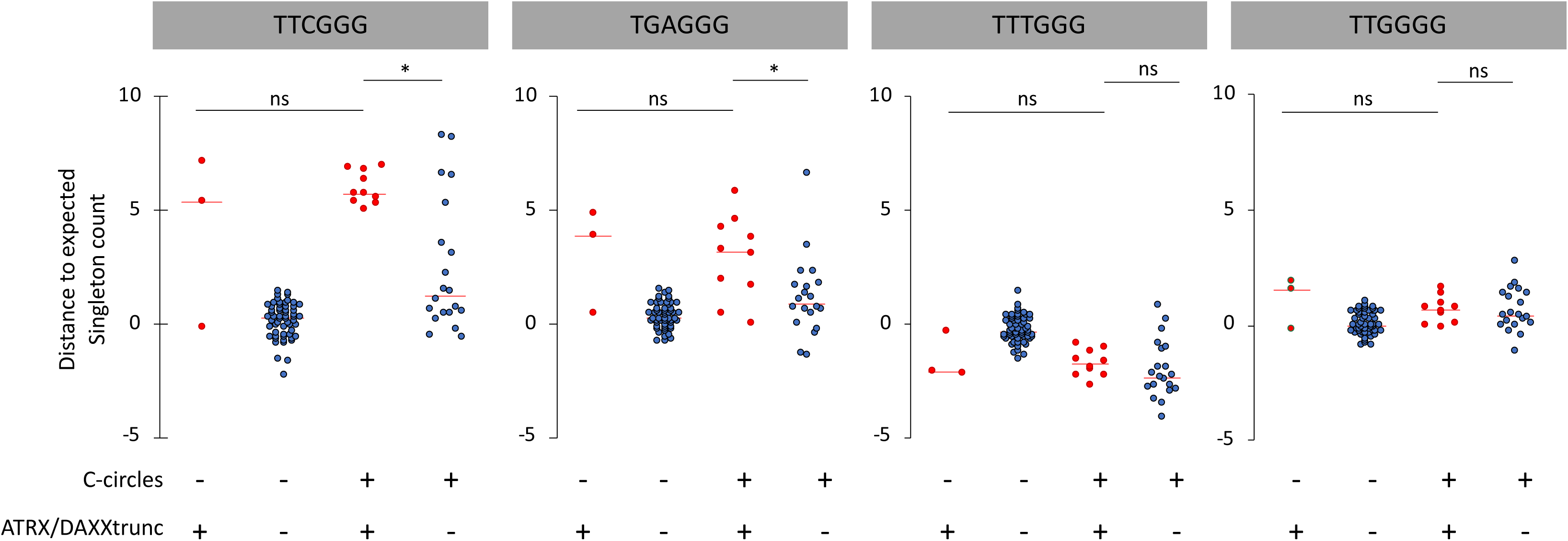
Distance to the expected singleton repeat count in C-circle negative and positive tumors with or without ATRX/DAXX^trunc^. The center (horizontal) red line of the scattergrams is the median. * *p* < 0.05, ns (non-significant); Kolmogorov-Smirnov tests.

ATRX is a SWI/SNF-like chromatin remodeling protein that binds to G-rich tandem repeats^11^. Once recruited, ATRX cooperates with histone chaperone DAXX in replication-independent deposition of the histone variant H3.3^12^. Loss of ATRX/DAXX function causes defects in multiple cellular processes, including defective sister chromatid cohesion and telomere dysfunction but is not sufficient by itself to induce ALT^13^. Our data suggest that ATRX/DAXX^trunc^ (rather than the ALT mechanism *per se*) could also play a role in TVR distribution. This observation could provide additional insights regarding the mechanisms involved in ALT promotion in the context of ATRX/DAXX^trunc^ in specific tumor types. TVR are associated with genomic instability, and could facilitate telomere spatial reconfiguration and affect telomere binding affinity^14–16^. Although histones are non-specific DNA binding proteins, nucleosome formation could potentially be influenced by telomeric DNA sequences and their structural properties^17^.

## A new classifier to identify ALT tumors in the PCAWG cohort

The international effort that enabled release of the PCAWG data set has provided a major new resource for seeking new insights into cancer, including the ALT pathway, which represents a potentially valuable, currently unexploited, target for anti-cancer therapies. Understanding more precisely how telomeres are maintained in cancer will shed light on replicative immortality and telomeric DNA damage response mechanisms. To develop a classifier, based on CCA data as an indicator of ALT, we searched for a signature associated with CCA status within the genomic features provided by the Telomere Hunter WGS tool^3^. Of the eight features used by Sieverling *et al.* in their random forest classifier, five were retained after Akaike information criterion stepwise regression analysis: telomere content (tumor/control log2 ratio) and the distance of TTTGGG, TTCGGG, TTGGGG and GTAGGG singletons from their expected occurrence. A classifier was then built from these five features, which permitted definition of two groups as “high-probability” and “low-probability”, using different combinations of the PanNET and melanoma datasets for training and testing. A 100% success rate was achievable when considering only one tumor type (i.e., using PanNET or melanoma for both the learning and the validation sets). Our final classifier, built using the whole cohort of patients with CCA data, has an accuracy of 93.86% (93.55% specificity and 93.98% sensitivity). This underlines the need for caution when extrapolating a classifier built on a specific tumor type to others. When applied to the 2,497 PCAWG patients, our classifier identified 200 tumors with high probability of ALT (Supplementary data 1), and their reported distribution across different histological types (leiomyosarcomas 73%, osteosarcomas 63%, liposarcomas 53% and low grade gliomas 29%) is consistent with previous reports^18^. In contrast, 27% of ATRX/DAXX^trunc^ tumors (n=17/64) were classified as ALT-low probability, and 6% of TERT^mod^ tumors (n=15/269) as ALT-high probability (Figure 1).

## Conclusion

The Sieverling *et al.* score is an ATRX/DAXX^trunc^ *vs.* TERT^mod^ classifier and is unsuitable for determining probability of ALT. The view that ATRX/DAXX loss is essentially equivalent to the presence of ALT activity may apply only to specific types of tumors. Adoption of this view as a generalisation across cancer types may divert attention from the need to identify alternative molecular actors involved in TMM. The classifier based on telomeric content and TVR distributions we provide here is more accurate as judged by C-circles, a hallmark of ALT, for PanNETs and melanomas, the tumor types on which it was trained. Although its predictions for other tumor types are consistent with the reported prevalence of ALT, this classifier should be applied with caution, given the apparent tumor-type specificity of TVR distribution.

## Data availability

The data used herein were used and referenced in published works by Sieverling *et al.*^3^ and Lee *et al.*^4^

## Contributions

A.N. and R.R.R. designed the study; A.N. performed the analysis.; A.N. and R.R.R. wrote the manuscript.

## Acknowledgements

Thanks to Prof. Hilda Pickett, Associate Prof. Karen MacKenzie and Yangxiu Wu for helpful comments and critical reading of the manuscript. A.N. is supported by a Doc.Mobility fellowship from the Swiss National Science Foundation.

---

**Supplementary figure 1.**
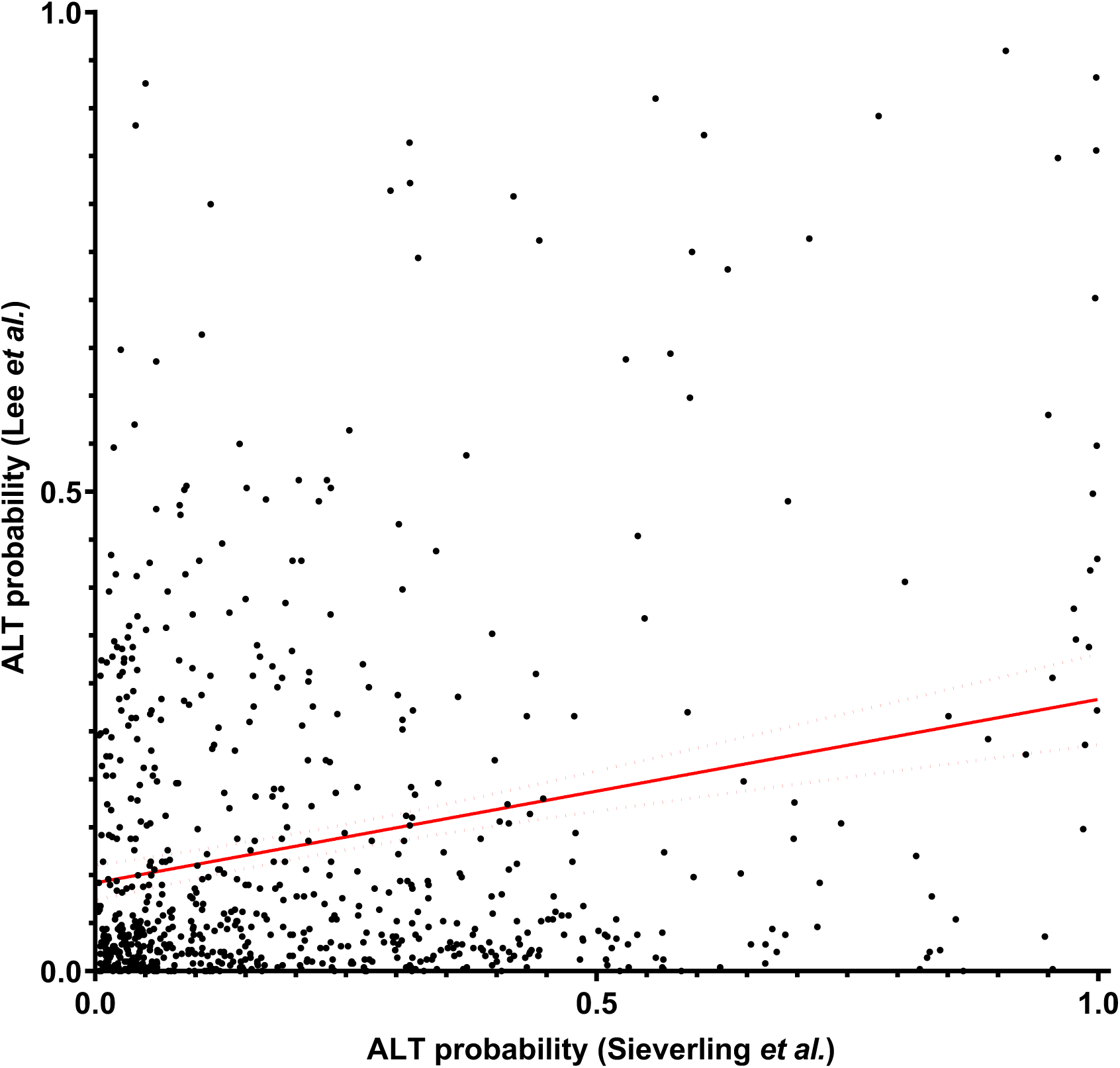
Correlation between Sieverling *et al.*^3^ and Lee *et al.*^4^ ALT-probability scores. Spearman *r*=0.101 [95%CI 0.025-0.17].

